# Dynamic changes to signal allocation rules in response to variable social environments in house mice

**DOI:** 10.1101/2022.01.28.478242

**Authors:** Caitlin H Miller, Matthew F Hillock, Jay Yang, Brandon Carlson-Clarke, Klaudio Haxhillari, Annie Y Lee, Melissa R Warden, Michael J Sheehan

**Author notes:** Authors for Correspondence: Caitlin H Miller, Michael J Sheehan.

## Abstract

Male house mice use metabolically costly urine marks in intrasexual competition and mate attraction. Given the high costs of signaling and the depletable nature of urine reserves, males should dynamically modulate signal allocation as the social landscape is updated with new information. We investigate which aspects of male urine marking behavior are static or dynamic in light of changing social environments. To do this, we use thermal imaging to capture spatiotemporal data of urine deposition decisions. This novel method reveals fine-scale variation in urinary motor patterns in response to competition and social odors. Males demonstrate striking winner-loser effects in both the total allocation effort and temporal dynamics of scent marking. We find that competitive experience primes key temporal features of signal allocation and modulates responses to familiar and unfamiliar male scents. Males adjust their signaling effort, mark latency, and scent mark rhythm, depending on the scent identities present in the environment. Winners dramatically increase marking effort toward unfamiliar compared to familiar male scent, consistent with a ‘dear enemy’ effect. Losers, in contrast, greatly reduce marking to unfamiliar scent but increase marking effort to the scent of their familiar rival, consistent with a ‘nasty neighbor’ effect. Counter to the high lability of many features, the initial signal investment pattern influences allocation decisions days later, revealing the possibility of alternative scent mark strategies among competitive males. Thus, different features of urine mark signal allocation vary in responsiveness to fluctuating social landscapes, suggesting there are multiple distinct behavioral modules underlying marking behavior.

## Introduction

Animals adjust their signaling behavior in response to recent experience and social context. Signalers may change not only the frequency of signaling behavior, but also when, where, and how they signal in response to changing social and physical environments (Hobson, 2020; Pasch et al., 2017; Patricelli & Hebets, 2016; Rauber & Manser, 2018; Sethi et al., 2019; Sullivan-Beckers & Hebets, 2014). Changes in allocation effort may allow individuals to take advantage of signaling opportunities or avoid unprofitable signal investment (Alberts, 1992; Ferkin, 2015; Gosling, 1982; Hurst & Beynon, 2004; Johnstone, 1996). In house mice (*Mus musculus domesticus*), males use metabolically costly urine marks to mediate intrasexual competition and mate attraction. The abundance, spatial distribution, and chemical composition of urine marks contain information about a male’s competitive status and identity (Desjardins et al., 1973; Drickamer, 2001; Ferkin, 2019; Gosling et al., 2000; Hurst, 1990; Hurst et al., 2001, 2005; Kaur et al., 2014; Lee et al., 2017; Nelson et al., 2015; Nevison et al., 2000). Previous studies have shown that the total level of urine deposition is modulated by social dominance (Desjardins et al., 1973; Drickamer, 2001; Hurst et al., 2005) and the presence of social odors in the environment (Hurst, 1990; Hurst et al., 2001; Kaur et al., 2014; Nevison et al., 2000). While urine marks convey rich social information, they are also directly depletable. Just as a car runs out of fuel, animals have a limited supply of urine to allocate toward signaling at any given moment. Thus, urine marks pose a unique set of production constraints that are not observed in other well-studied signaling systems, such as songs or visual displays (Cooper & Goller, 2004; Gil & Gahr, 2002; Laidre & Johnstone, 2013; Ruppé et al., 2015). We explore the flexibility of signal allocation decisions, both on a moment-to-moment timescale as well as over the course of days. Understanding how individuals integrate social information to allocate depletable and costly signals has the potential to reveal fundamental features of complex decision-making processes.

Male social relationships are shaped by competition and familiarity with conspecifics in house mice (Anderson & Hill, 1965; Crowcroft & Rowe, 1963; Desjardins et al., 1973; Harrington, 1976; Koolhaas et al., 2013; Mackintosh, 1970; Poole & Morgan, 1976; Wolff, 1985). Urine marking mediates some of these relationships by allowing assessment and recognition of individuals (Drickamer, 2001; Hurst et al., 2001, 2005; Kaur et al., 2014; Nevison et al., 2000). Both stimulus familiarity and aggressive contests independently have strong effects on male urine marking (Arakawa, Arakawa, et al., 2008; Arakawa, Blanchard, et al., 2008; Desjardins et al., 1973; Drickamer, 2001; Jones & Nowell, 1973), however it remains poorly understood how the two interact. One possibility is male mice respond indiscriminately to the urine of other males (i.e. non-self) regardless of their experience with said individuals. Another possibility is males respond differentially to the urine of males with whom they have established relationships versus unfamiliar novel males. In many territorial species, familiar neighbors reduce aggressive behaviors and signaling effort toward familiar individuals known as the “dear enemy” effect (Booksmythe et al., 2010; Briefer et al., 2008; Christensen & Radford, 2018; Tumulty & Bee, 2021; Zorzal et al., 2021). The dear enemy effect is well-documented across vertebrate species, and is thought to lessen the costs of territorial defense (Tumulty, 2018). Alternatively, marking may be influenced by a “nasty neighbor” effect (Christensen & Radford, 2018; Goll et al., 2017; Jin et al., 2021; Müller & Manser, 2007), where territorial males increase their signaling effort toward familiar neighbors. Given the high costs and depletable nature of urine marks, males should dynamically modulate signal allocation as the landscape is updated with new social information. The present study aims to shed some light on these decision rules by exploring how established competitive relationships and identity information influence male signal allocation decisions across social and scent-marked environments.

We investigate how males shift their signal allocation in response to an aggressive contest, to the presence of a familiar male competitor, and to the presence of urine scent-marks of differing male identities. The objectives of this study were to: (1) establish thermal imaging as a novel method for measuring scent marking, (2) explore how competitive experience alters marking behavior, and (3) test the hypothesis that familiarity is important for signal allocation decisions. To do this, we developed a 4-day trial design in which 31 pairs of age and weight-matched breeding males of distinct wild-derived strains were paired as competitors and presented a series of social and scent-marked trials (Figure 1A). Two wild-derived partially inbred house mouse strains were used to examine scent marking behaviors, and ensured that males of each pair smelled distinct from one another (Kaur et al., 2014; Sheehan et al., 2016, 2019). On the first day, paired males were placed in an arena separated by a mesh barrier (Figure 1A). Paired males could see, hear, and smell each other but were limited to minimal physical contact through the mesh. The mesh barrier was subsequently removed, and males engaged in an aggressive contest or “fight trial” (Figure 1A). Based on the total aggressive behaviors performed by each male in the fight trial, males were easily and unambiguously classified as winners or losers (Figure S1). On the second day, each male was placed in an empty open field arena without any stimuli aside from the arena itself (Figure 1A). On the third day, males were placed back into the mesh arena with the same male competitor encountered on the first day (Figure 1A). Finally, on the fourth day, each male was exposed to one of four urine mark treatments. Each treatment contained aliquoted male urine of three possible identities (self, familiar male, or unfamiliar male) in two spatially distinct scent-marked zones (Figure 1A). The four treatment types span a range of scent mark combinations (self-self, self-familiar, self-unfamiliar, familiar-unfamiliar), in which the familiar stimulus is the urine of a male’s paired competitor (Figure 1A). Urine marking behavior and space use data were collected for each male across urine marking assays while aggression was scored in the fight trials (Figure 1B-C; Figure S1).

**Figure 1.**
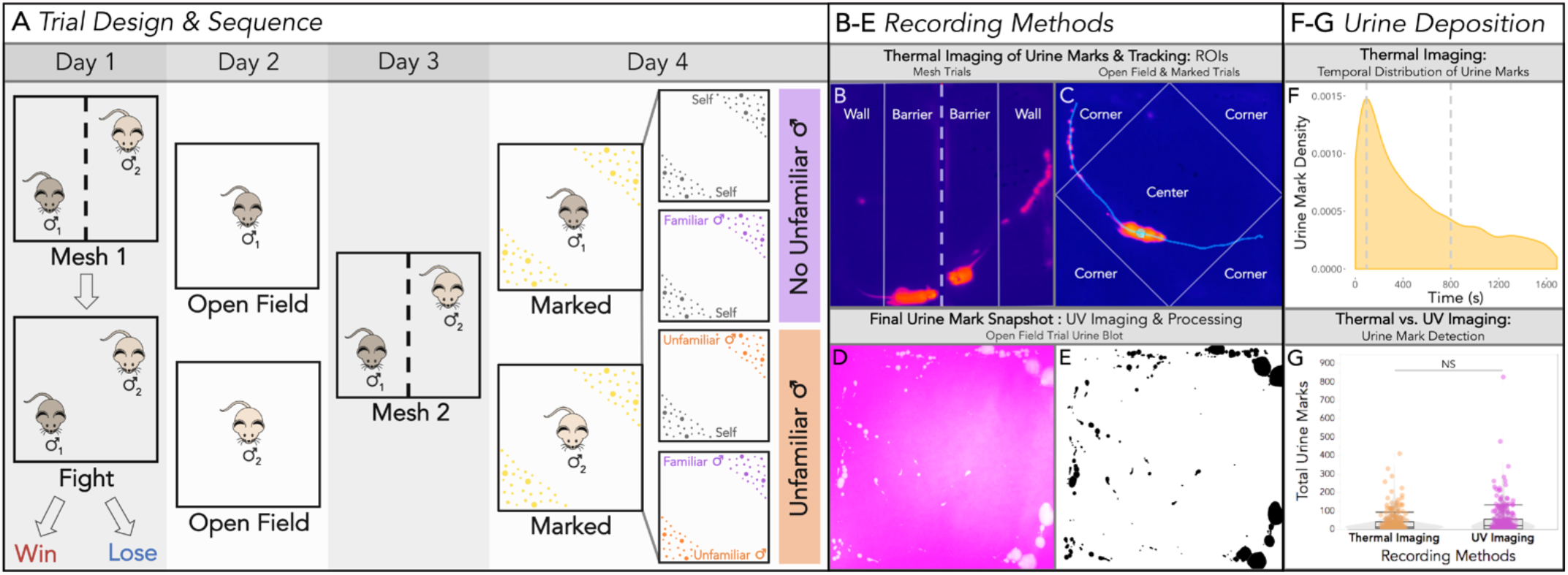
Experimental design and recording methods. **(A)** Trial design for a given pair of mice. All trials were 30 minutes long. Day 1: age and weight-matched males of distinct wild-derived strains were paired as competitors and placed into an arena and separated by a mesh barrier (dashed line). The mesh barrier was removed and males entered into an aggressive contest (fight trial) concluding in winning or losing males. Day 2: each male was placed into an empty open field arena. Day 3: males were placed back into the mesh arena with the same (familiar) male competitor from the first trial day. Day 4: each male was exposed to one of 4 possible treatments of aliquoted male urine of 3 possible identities (self, familiar and unfamiliar) into two urine-marked zones. The 4 treatment groups: self-self, self-familiar male, self-unfamiliar male, familiar male-unfamiliar male. The familiar male stimulus is the urine of a male’s paired competitor (present in mesh and fight trials). **(B)** Thermal recording snapshot of a mesh trial (Days 1 & 3) depicting the regions of interest (ROIs: Wall vs. Barrier) used to score urine mark deposition and track space use. The dashed line indicates the mesh barrier separating the two males. The solid lines depict the ROIs each male can traverse on their side of the barrier. Urine marks are hot (orange-pink: close to the body temperature) on a cool (dark blue) ambient substrate (filter paper) temperature. The first and last minute of each scent-marking trial was trimmed prior to analysis (total analyzed trial length: 28 minutes) to minimize detection of startle-based urination events caused by placement of mice into arenas and any jostling caused during trial set-up and take-down. **(C)** Thermal snapshot of open field (Day 2) and urine-marked (Day 4) trials with the ROIs used for scoring (Corners vs. Center) indicated with solid lines. An example track of the mouse’s trajectory two seconds before and after its current location is shown (light turquoise) with the mouse body’s centroid indicated with a circle. Recently deposited urine marks (hot pink) align with the past track of the focal mouse. **(D-E)** Example final urine blot (on filter paper) of an open field trial imaged under UV light with urine marks fluorescing brightly (D), and the processed inverted urine blot image with urine marks shown in black (E). **(F)** Density plot depicting the temporal distribution of all thermally detected urine marks across all trials. **(G)** The total number of urine marks detected across trials using novel thermal imaging and standard UV blot imaging recording methods. A linear mixed model (LMM) was used to model the relationship between recording method and total urine marks detected. An analysis of variance (ANOVA) was used to test for the overall effect of recording method (significance code: NS p>0.05).

## Results

### Thermal imaging reveals the spatiotemporal dynamics of scent marking in real time

To best study urine allocation decisions in mice, we need to measure real-time spatial and temporal patterns of scent mark deposition events. Mouse urine marking has been previously studied by capturing final snapshots of marking patterns at the end of a trial, providing a cumulative output of behavior (Desjardins et al., 1973; Kaur et al., 2014). More recently, methods have been developed for tracking temporal changes in urine output by filming under UV light or a moving paper floor (Hou et al., 2016; Keller et al., 2018). Here, we used thermal imaging as an unobtrusive method for capturing in real time both the spatial and temporal allocation of urine marks by male house mice (Figure 1). Urine leaves the body hot (close to body temperature) and quickly cools to below the ambient substrate temperature, providing a distinctive thermal signature. In this study we examine both the *where* and the *when* of urine mark deposition. Trials were performed on filter paper to present urine stimuli and to generate informative summary images for each trial by imaging the urine blots under UV light (Figure 1D-E). This further allows for comparison of thermal recording with a traditional urine detection method.

Using thermal imaging we recorded a total of 9,314 urine deposition events across trials, and we explored the temporal distribution of these depositions. We observe an initial spike in urine deposition with a peak of activity at ∼100s, followed by an exponential decline for the remainder of the trial length (Figure 1F). The majority (77%) of marks are deposited within the first 15 minutes (800s) of the trial, suggesting males rapidly scent mark upon entering an environment (Figure 1F). Thermal imaging focuses on urine *deposition*, as marks are scored by the distinct thermal profile of urine as it is deposited. UV light imaging cannot distinguish between deposition and *distribution* events, as urine is further distributed by males walking through their own marks and tracking urine with their paws and tail. However, UV imaging can also undercount urine marks when depositions occur in the same location. UV light imaging at times detects more spots of urine than the true number of marks detected by thermal imaging across trials, though not significantly (*F*_1,430_ = 0.0034, *p* = 0.95; Figure 1G). The two detection methods are also highly correlated (Figure S2), justifying the use of thermal imaging to examine how temporal urine allocation varies across contexts.

### Competitive experience and initial signal investment shape urine mark allocation

Competitive social encounters can have a range of important consequences on the behavior and physiology of individuals (Harrison et al., 2018; Hsu & Wolf, 1999; Li et al., 2014; Milewski et al., 2022; Rose et al., 2017; Thomas et al., 2015; Tibbetts & Crocker, 2014). How individuals respond to contest outcomes is often dependent on their assessment of their own resource holding potential, which is often further updated through experience (Arnott & Elwood, 2009; Briffa & Elwood, 2009; Enquist & Leimar, 1983; Humphries et al., 2006). Signals play a key role in such encounters as they can convey information about the competitive ability of individuals (Kodric-Brown & Brown, 1984; Ligon & McGraw, 2016; Számadó, 2017; Tibbetts & Izzo, 2010). In house mice, initial scent mark investment by males has been shown to contain information about their competitive ability (Drickamer, 2001). We therefore predicted that (1) higher-marking males would be more likely to win aggressive contests, (2) winners would increase while losers would decrease signaling after a contest, and (3) the changes in signaling after a contest may depend upon their initial signaling strategies. We thus compared how males allocate urine marks in the presence of a competitor *before* and *after* a fight by comparing the mesh trials on days 1 and 3 of the trial series (Figure 1A).

We classified winners and losers based on the total number of aggressive behaviors performed by each male in the fight trial (Figure S1). In all contest pairings the fight outcome was overwhelming clear (Figure S1A), with winning males performing significantly more aggressive behaviors than losing males (*t*_1,31_= -12.6, *p* = 1.09e-13; Figure S1B). Fight outcome has a strong effect on total urine marks (*F*_1,68_ = 10, *p* = 0.0021), and there is a significant interaction between fight outcome and trial number (*F*_1,60_ = 12, *p* = 0.0011). Before the fight (Mesh 1), the to-be winners include more high-marking individuals than the to-be losers, however, the two groups did not differ significantly (*t*_1,112_ = -0.69, *p* = 0.88; Figure 2A). Initial signal investment may carry some information about competitive ability (Drickamer, 2001), but this did not predict fight outcome between age and weight-matched males in our trials. Post-fight (Mesh 2), the total urine marks deposited by winners is significantly higher than losers (*t*_1,112_ = -4.6, *p* = 0.0001; Figure 2A). Similar to previous studies (Arakawa, Arakawa, et al., 2008; Arakawa, Blanchard, et al., 2008; Desjardins et al., 1973; Drickamer, 2001), this relationship appears to be largely driven by a decrease in urine marking among losing males (*t*_1,61_ = 3.3, *p* = 0.0059; Figure 2A,B).

**Figure 2.**
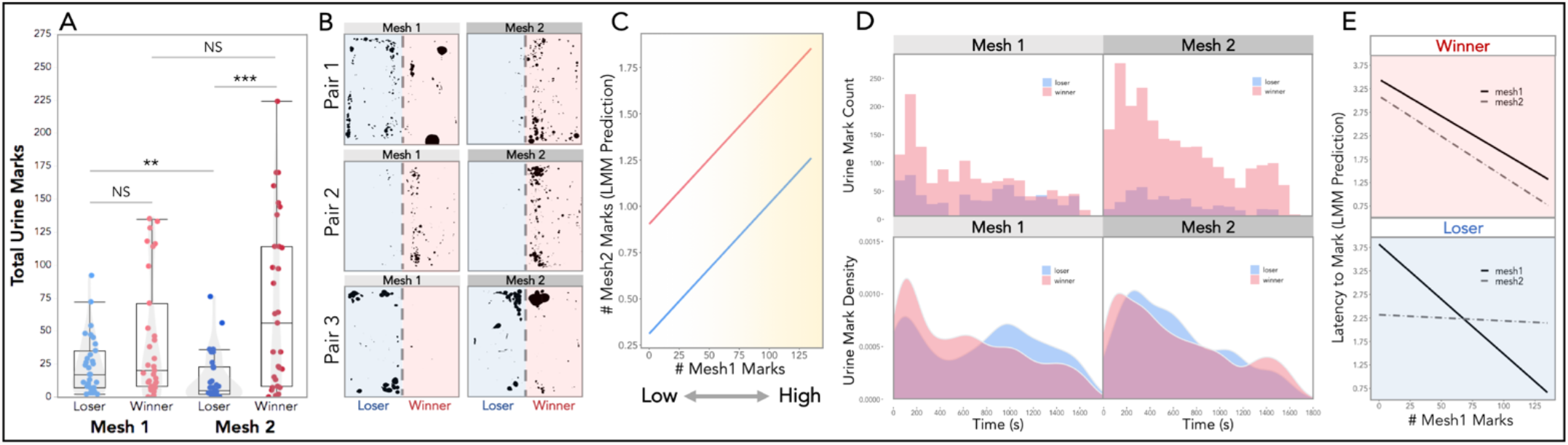
Male urine mark allocation in response to social competition across mesh trials. **(A)** Total urine marks deposited in Mesh 1 (pre-fight) and Mesh 2 (post-fight) by losers and winners. **(B)** Mesh trial urine blots of three paired male competitors (winner and loser) pre- and post-fight. **(C)** Linear mixed model (LMM) prediction of the total number of Mesh 2 marks (log-transformed) given fight outcome (winner=red; loser=blue) and initial signal investment (total number of Mesh 1 marks). **(D)** Histograms (top) of the temporal distribution of urine marks deposited by winners (red) and losers (blue) in Mesh 1 (pre-fight) and Mesh 2 (post-fight). Density plots (bottom) depict the density of urine mark deposition events over the 30-minute Mesh trials, distinguished by fight outcome (winner=red; loser=blue). **(E)** Linear mixed model (LMM) prediction of the latency to mark in both Mesh trials given the fight outcome (winner=red; loser=blue) and initial signal investment (total number of Mesh 1 marks). **(A**,**C**,**E)** Linear mixed models (LMMs) were used to model relationships, and analyses of variance (ANOVAs) were used to test for overall effects (significance codes: NS p>0.05, * p<0.05; ** p<0.01, *** p<0.001). Dependent variables were logarithmically transformed to meet assumptions for model residuals.

We next assessed the role of initial signal investment (# Mesh 1 marks) and fight outcome on subsequent allocation patterns (Figure 2C). Given prior research, we expected some males would mark highly, lose the fight, and then suppress their marking behavior in the later trial (Desjardins et al., 1973). Instead, we found that the importance of initial signal investment is more generalizable than this anticipated loser effect (Figure 2C). How much an individual marked in the first mesh trial has a strong effect on the urine mark allocation in the second mesh trial (*F*_1,59_ = 9.2, *p* = 0.0036; Figure 2C). In other words, if you start off a low-marking individual you remain *relatively* low-marking, regardless of the fight outcome. Accordingly, both high-marking losers and low-marking winners are observed (Pair 3: Figure 2B). The pronounced winner-loser effects of urine mark allocation are therefore strongly modulated by initial signal investment.

Losing has a notable effect not only the on number of marks, but also *where* individuals place those marks in the arena (Figure S3). On the second mesh trial, losers allocate their marks differently in space (at the wall vs. barrier) depending on whether they started off as high or low-marking (Figure S3A,B), suggesting that losers may be altering signaling strategies in addition to total signaling effort. Space use, on the other hand, does not differ (Figure S3C). All individuals spend more time in the social region of the arena (barrier) regardless of fight outcome (Figure S3C). Surprisingly, where males spend time does not correlate with where they mark (Figure S3D), indicating males are not simply marking where they spend time but are specifically allocating their urine marks.

### Social experience influences the temporal dynamics of scent mark allocation

In addition to the total number of urine marks, mice may alter the relative timing of urine mark deposition, such that marks may be more spaced out or clustered in time. The relative timing of urine deposition provides novel information on the instantaneous rates of signaling, revealing how mice choose to spend their urine reserves. A slow and regular mark deposition strategy is distinct from marking in rapid bursts upon the detection of a stimulus or entry into an environment. In more natural contexts it is very plausible that rapid bursts of marking facilitate competitive signaling, as males are traversing larger distances and navigating more complex social environments.

We inspected the distribution of urine mark deposition for winners and losers across mesh trials (Figure 2D). In the first trial, winners and losers display an initial peak in urine deposition at ∼100 seconds (Figure 2D). Notably, the losers-to-be have a distinct second peak much later in the trial (∼1000 seconds; Figure 2D). In the second mesh trial the effect of fight outcome is clear, with winners marking considerably more and losers less (Figure 2D). The density curves, however, reveal that both winners and losers allocate their marks earlier in the trial followed by a sustained decline in deposition events (Figure 2D). This shift in urine deposition regardless of fight outcome suggests a general priming effect of social competition on the temporal allocation of urine marks.

The timing of a male’s first scent mark changes with fight outcome (Figure 2E). Mark latency is strongly influenced by initial signal investment (*F*_1,59_ = 39, *p* = 4.4e-08). Similarly, trial order has a clear effect on the latency to mark (*F*_1,58_ = 10, *p* = 0.0021), while fight outcome does not (*F*_1,57_ = 0.24, *p* = 0.62). The three-way interaction between trial order, fight outcome, and initial mark investment significantly effects mark latency (*F*_1,58_ =10, *p* = 0.0021; Figure 2E). For both winners and losers, low-marking males are slower to mark than high-marking males, characterizing a low and slow marking pattern on the first day. Conversely, high-marking individuals typically mark rapidly upon entering the arena on the first day, representing a high and fast marking pattern. Across the two trial days, winners speed up their mark latency, though this effect is scaled to their initial mark investment (Figure 2E). Initially high-marking losers have an increased latency after losing (i.e. they are slower to mark), but individuals who marked infrequently on the first trial decrease their latency, demonstrating complex shifts in signaling behavior dependent on initial state. (Figure 2E).

We next examined the timing and rhythm of urine marking across entire mesh trials. Event plots depicting the timing and frequency of marks deposited for two male pairs are shown (Figure 3A). The winning males increase the front-loading of urine deposition in the second trial, whereas the losers decrease overall mark deposition but still mark relatively quickly in the second trial (Figure 3A). Examining the timing of marks across the trial reveals unanticipated patterns as well. The intervals between marks differ noticeably between the first and second mesh trials, particularly when marks are made in close sequence to each other. In the first trial, the mark sequences have larger intervals, producing chains of marking events (Figure 3A & S4). Whereas in the second mesh trial the mark sequences are compressed into shorter intervals, creating bursts of marking events (Figure 3A). To examine this relationship further, we inspected the distribution of inter-mark intervals (IMIs) among winners and losers for both trials (Figure 3B). On the first trial the median IMI of winners and losers is similar (Figure 3B). However, on the second trial, the median IMI for losers increases, driven by the fact that fewer marks are being made overall. While IMIs have a wide range, the most frequent mark intervals are under 3 seconds (Figure 3B & S4). In the first trial, the most frequent IMIs are less than 3 seconds for winners and losers, though winners have a lower median mark value. (Figure 3B). In the second trial, there is a clear peak of IMIs of less than 1 second for both winners and losers (Figure 3B). This suggests that both winners and losers are producing mark “chains” on the first trial and “bursts” of marks on the second. The overall median IMI interval is unchanged for winners but increases notably in losers.

**Figure 3.**
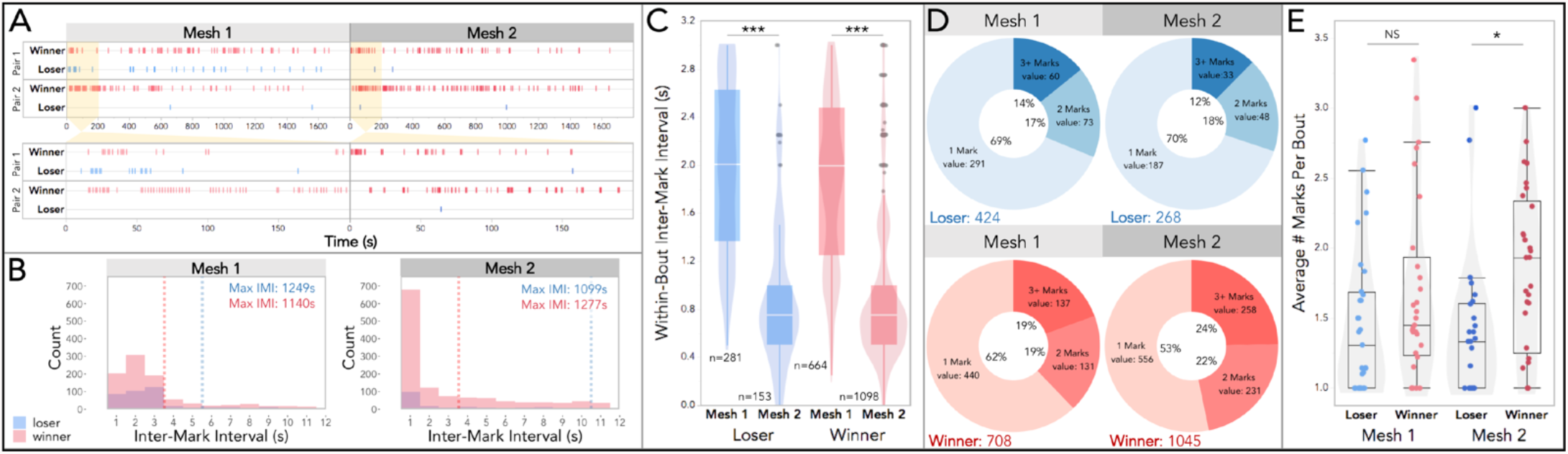
Temporal dynamics of urine mark allocation across mesh trials. **(A)** Example data event plots depicting the urine marking of two pairs of male competitors over the course of both mesh trials for the entire trial duration (top) and a zoomed-in view of the first 200s (bottom). **(B)** Histograms of the inter-mark intervals (IMIs) for winners and losers in both mesh trials. Median values are indicated with dashed lines. The range of IMIs extends to nearly the full trial length (only the first 12s is shown). The maximum values are reported in the top right corner. Mesh 1: 65% of all IMIs are shown (< 12s), 57% of loser IMIs and 69% of winner IMIs. Mesh 2: 68% of all IMIs are shown (< 12s), 51% of loser IMIs and 72% of winner IMIs. **(C)** Box and violin plots of within-bout IMIs by fight outcome and mesh trial. **(D)** Donut plots by trial and fight outcome depicting the proportion of bouts composed of: 1 mark, 2 marks or 3+ marks. Mark totals in the bottom left-hand corner. **(E)** Boxplot of the average number of marks per bout by fight outcome and mesh trial. **(D**,**E)** Linear mixed models (LMMs) were used to model relationships, analyses of variance (ANOVAs) were used to test for overall effects, and post hoc pairwise comparisons were performed using the *emmeans* package (significance codes: NS p>0.05, * p<0.05; ** p<0.01, *** p<0.001).

To explore this shift in temporal dynamics *within* urine mark sequences, we classified marks that occur in inter-mark intervals of less than 3 seconds as marking bouts (Figure S4). Bouts can thus consist of a single mark or a series of marks. We then examined the variation in IMIs within urine mark bouts (Figure 3C). As expected, trial has a strong effect on within-bout IMIs (i.e. IMIs for bouts with 2+ marks, *F*_1,428_ =304, *p* = 2.0e-16) while fight outcome does not (*F*_1,46_ = 0.079, *p* = 0.78; Figure 3C). Thus, marking events within bouts are more rapid in the second trial, regardless of whether they won or lost, indicating that competitive experience may prime a particular marking motor pattern, regardless of fight outcome.

We next inspected whether bouts are composed of 1 mark, 2 marks or 3+ marks (Figure 3D). On the first trial losers have more single-mark bouts and winners have more multi-mark bouts (Figure 3D). This relationship becomes even more stark in the second mesh trial, as losers decrease the overall number of marks across mesh trials, but the bout composition remains very similar (Figure 3D). Winners, on the other hand, increase the number of marks *and* alter their bout composition to include more multi-mark bouts (Figure 3D). To further explore bout composition, we compared the average number of marks per bout by fight outcome and trial (Figure 3E). In contrast to the IMIs of urine mark bouts, bout composition is strongly affected by fight outcome (*F*_1,58_ = 10, *p* = 0.0022; Figure 3E). On the second trial, winners have a significantly higher average number of marks per bout than losers (*t*_1,111_ = -3.0, *p* = 0.012; Figure 3E). This dataset reveals striking patterns of signaling behavior in male house mice that would have otherwise gone unnoticed without temporal data possible from thermal imaging.

### Dominance and familiarity interact to shape countermarking behavioral dynamics

Given that males dynamically adjust marking behavior in response to social competition, we next explored allocation decisions toward the scent marks of other males. We were especially interested in whether males use knowledge of a recent competitor’s identity in their signaling decisions, as males will competitively counter-mark to (i.e. mark over) the urine marks of other males (Hurst & Beynon, 2004; Kaur et al., 2014). While it is well-established that males alter marking behavior in response to fight outcome (Desjardins et al., 1973; Drickamer, 2001; Hurst, 1990) and can finely discriminate urine identities (Hurst et al., 2001; Kaur et al., 2014), we have a limited understanding of how males implement this information in a competitive marking context. Do males adjust their scent marking behavior depending on their relationship to a male competitor? What role does familiarity play in signal allocation dynamics? We hypothesized that fight outcome would shape urine marking behavior even in the absence of a male competitor, and that male identity in urine-marked trials would strongly govern signal allocation decisions.

To address these questions, we compared two trial types within the trial series in which no conspecifics were present: open field trials (OFTs) and urine-marked trials (Figure 1A). The OFT contained no stimuli, while the urine-marked trials each contained two spatially distinct urine-marked zones of specific identities: their own urine (self: S), familiar male (FM) competitor urine, and/or unfamiliar male (UM) urine (Figure 4B). UM urine was collected from a different strain (C57BL/6J) and pooled to ensure a distinct urine profile with no individual-specific effects on the UM stimulus. We examined responses to an empty arena (OFT) and to the four different urine stimulus sets: S-S, S-FM, S-UM and FM-UM (Figure 1A) by fight outcome and initial signal investment. Trial type (*F*_4,76_ = 5.2, *p* = 0.00089), fight outcome (*F*_1,83_ = 27, *p* = 1.6e-06), and initial signal investment in the first trial (*F*_1,58_ = 32, *p* = 4.2e-07), all significantly effect marking behavior of males (Figure 4A). As does the two-way interaction between trial type and fight outcome (*F*_4,77_ = 5.8, *p* = 0.00041). Winners tend to mark more than losers, and losers typically mark lowly across treatment types. This pattern is further observed in the empty arena trials (*t*_1,100_ = -3.9, *p* = 0.0034; Figure 4A). Notably, winners and losers show opposite responses towards familiar versus unfamiliar urine. Treatments with only familiar urine (Figure 4A,B, purple: S-S and S-FM) exhibit comparable marking responses in winners and losers (Figure 4A). While it’s perhaps less surprising that winners and losers mark comparably lowly to their own urine (S-S; *t*_1,99_ = -0.83, *p* = 1.00), it is striking that winners and losers do not differ in their response to the S-FM treatment (*t*_1,105_ = -0.44, *p* = 1.00; Figure 4A). The opposite pattern is observed in the presence of unfamiliar urine. Winners mark significantly more than losers to both S-UM (*t*_1,108_ = -3.6, *p* = 0.0091) and FM-UM (*t*_1,109_ = -6.0, *p* < 0.0001) treatments (Figure 4A). In trials with two different urine identities, we had anticipated males would differentially allocate urine towards each marked corner, though we did not detect such differences in either winners or losers (Figure S5A). While scoring the trials, it became clear that the space was too small to delineate corner-based stimuli as distinct ROIs like we had originally planned. Especially since males frequently walk through corners while performing a scent-mark bout that extends across multiple ROIs. Furthermore, our results indicate males may respond to the most ‘extreme’ (or most salient) odor information in the environment within the spatial confines provided (Figures 4 & 5). However, space use patterns of losing males suggest that mice are detecting differences in the scent marks (Figure S5B). Losers spend less time in the center ROI compared to winners (*t*_1,230_ = -3.7, p = 0.0069), and losers spend less time in UM-marked corners relative to empty ones (*t*_1,199_ = -3.5, p = 0.0012) (Figures 1C & S5B).

**Figure 4.**
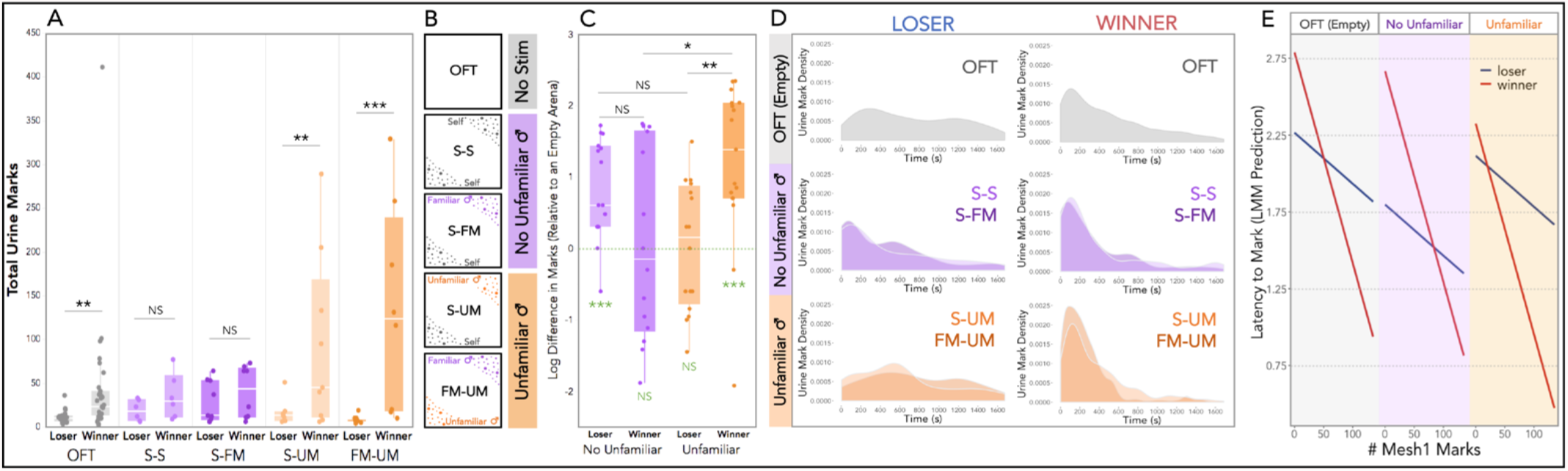
Urine mark allocation across scent-marked contexts in response. **(A)** Total urine marks deposited by winning and losing males in the open field trial (OFT) and the four urine-marked treatments: self-self (S-S), self-familiar male (S-FM), self-unfamiliar male (S-UM) and familiar male-unfamiliar male (FM-UM). All males experienced the OFT. Males experienced only one of the four urine-marked treatments. **(B)** Schematic of the urine stimulus components for the OFT and urine-marked treatments. OFTs are “no stimulus” trials (grey), S-S and S-FM have “no unfamiliar male” urine present (purple), and S-UM and FM-UM trials have “unfamiliar male” urine present (orange). **(C)** The difference in total marks deposited by males in the Marked trials relative to the empty OFTs (logarithmically transformed). Urine-marked treatments are grouped as “no unfamiliar male” urine (purple: S-S and S-FM) and “unfamiliar male” urine (orange: S-UM and FM-UM). Post hoc pairwise comparison significance values indicated at the top of boxplots. One-sample t-tests (deviation from 0) significance values are indicated on the bottom of the boxplots **(D)** Urine mark density plots of losing and winning males toward an empty arena (OFT), to trials with no unfamiliar male urine: S-S (light purple) and S-FM (dark purple), and to trials with unfamiliar male urine: S-UM (light orange) and FM-UM (dark orange). **(E)** Linear mixed model (LMM) prediction of the latency to mark in the OFT trials (gray), Marked trials with “No Unfamiliar” male urine (purple), and Marked trials with “Unfamiliar” male urine present (orange), given the fight outcome (winner=red; loser=blue) and initial signal investment (total number of Mesh 1 marks). **(A**,**C**,**E)** Linear mixed models (LMMs) were used to model relationships, analyses of variance (ANOVAs) were used to test for overall effects, and post hoc pairwise comparisons were performed using the *emmeans* package (significance codes: NS p>0.05, * p<0.05; ** p<0.01, *** p<0.001). Dependent variables were logarithmically transformed to meet assumptions for model residuals.

We observed very similar responses in the two treatments with unfamiliar urine present (S-UM and FM-UM) as well as the two treatments without unfamiliar urine (S-S and S-FM), so we collapsed these similar treatments (purple: no unfamiliar male, orange: familiar male) to further explore the role of familiarity and fight outcome on signal allocation (Figures 4B-E). We standardized the marking behavior of males by calculating the difference in marks made in an empty arena (OFT) relative to a scent-marked environment (Figure 4C). The interaction between fight outcome and familiarity strongly shapes marking behavior in scent-marked contexts (*F*_1,58_ = 13, *p* = 0.00054; Figure 4C). Winners increase the number of marks significantly more than losers in trials with unfamiliar urine present (*t*_1,58_ = -3.0, *p* = 0.0069), whereas winners and losers do not differ when familiar-only scent marks are present (*t*_1,58_ = 2.0, *p* = 0.17; Figure 4C). What is also apparent is the inverse response of winners and losers toward familiarity (Figure 4C). Winners mark relatively more in the presence of unfamiliar urine and lowly to familiar-only urine (*t*_1,58_ = -3.0, *p* = 0.014), while losers mark lowly in the presence of unfamiliar urine and highly to familiar-only urine (*t*_1,58_ = 2.2, *p* = 0.12; Figure 4C). Notably, losers in the familiar-only treatment (*t*_1,14_ = 4.5, *p* = 0.00048) and winners in the unfamiliar treatments (*t*_1,16_ = 4.4, *p* = 0.00041) deviate significantly from zero, while their opposing treatments do not (Figure 4C).

### Temporal variation in signal allocation during countermarking

The timing of signal allocation in scent-marked environments was examined (Figure 4D). We first looked at the distribution of deposition events across trials (Figure 4D). In trials with no urine stimulus (OFTs), winners allocate marks early in the trial (peak density ∼100s), while losers mark less with a later peak at ∼250s (Figure 4D). In contrast, winners and losers have remarkably similar density curves for urine-marked trials containing familiar-only urine (purple: S-S and S-FM) (Figure 4D). For S-S trials, both winning and losing male density curves display a single initial peak (∼100s), whereas the S-FM trials’ density curves reveal an earlier initial peak (∼70s) and a smaller second peak later in the trial (Figure 4D). In trials with unfamiliar urine, winners and losers differ dramatically. Winning males deposit large amounts of urine very quickly, creating a large initial spike in the density curves in both S-UM (light orange) and FM-UM (dark orange) treatments (Figure 4D). Losing males drop off and slow down their urine mark deposition considerably, generating density curves with small and delayed peaks (Figure 4D). The temporal distribution of urine marks is thus modulated by fight outcome and familiarity in scent-marked environments.

As the temporal dynamics of scent-marks were overlapping in trials with or without unfamiliar male urine, we collapsed these into treatment groups (Figure 4E). We further modeled the effects of treatment group, fight outcome, and initial signal investment, on the latency to mark (Figure 4E). Mark latency is significantly predicted by the number of marks made in the first mesh trial, i.e. the initial investment recorded 3 days earlier (*F*_1,57_ = 10, *p* = 0.0024; Figure 4E). For winners and losers, initially low-marking individuals are slower to mark, and initially high-marking individuals are faster to mark (Figure 4E). This relationship is most stark among winners, which exhibit steep slopes across treatment groups, while losers display more modest slopes (Figure 4E). The interaction between fight outcome and initial signal investment, however, is moderate (*F*_1,57_ = 3.7, *p* = 0.056 ; Figure 4E). The effect of fight outcome on mark latency is not significant (*F*_1,56_ = 1.3, *p* = 0.25 ; Figure 4E). Treatment group on the other hand, significantly effects the speed of marking response (*F*_1,75_ = 3.2, *p* = 0.048; Figure 4E). Losers mark most rapidly in familiar-only trials, and winners mark most rapidly in trials with unfamiliar urine (Figure 4E). The intersection points of the linear models for winners and losers reveal further insights. Winners transition to a more rapid marking response relative to losers differently across treatments groups depending on initial signal investment. In familiar-only trials, only the initially very high-marking (>85 marks) winners mark more quickly than losers, others are slower to mark. The opposite is true in trials with unfamiliar urine, in which even initially low-marking (>20 marks) winners mark more rapidly than losers (Figure 4E). This demonstrates that initial signal investment has long-term power for predicting marking behavior, including the temporal allocation of urine marks.

We next examined the timing and composition of marking bouts (Figure 5). More chain-like bouts are observed in OFTs, whereas more rapid bursts of urine marking are produced in scent-marked trials (Figures 5A, 5D & S5C). Therefore, over the 4-day trial series males mark increasingly in bursts, suggesting competitive experience shapes temporal features of signal allocation. To explore this further we looked at within-bout IMIs (Figure 5B). Both fight outcome (*F*_1,64_ = 6.2, *p* = 0.015) and treatment group (*F*_1,152_ = 40, *p* = 1.2e-14) significantly effect within-bout IMIs, with a modest interaction (*F*_1,154_ = 2.5, *p* = 0.083; Figure 5B). As expected, the within-bout IMIs are significantly longer in OFTs than either scent-marked treatment groups among winners or losers (Figure 5B). Winners, however, marked with similar rapid bursts (short IMIs) regardless of familiarity with the urine stimulus (Figure 5B). Conversely, losers tend to mark in bursts specifically during familiar-only trials (Figure 5B). This bout timing is most prominent in the S-FM trials (Figure 5A & S5C), which reveals losers distinctly allocate their urine marks based on the identity of urine marks in the environment. It is striking that, again, losers signal most conspicuously toward males who recently defeated them in a competitive contest.

**Figure 5.**
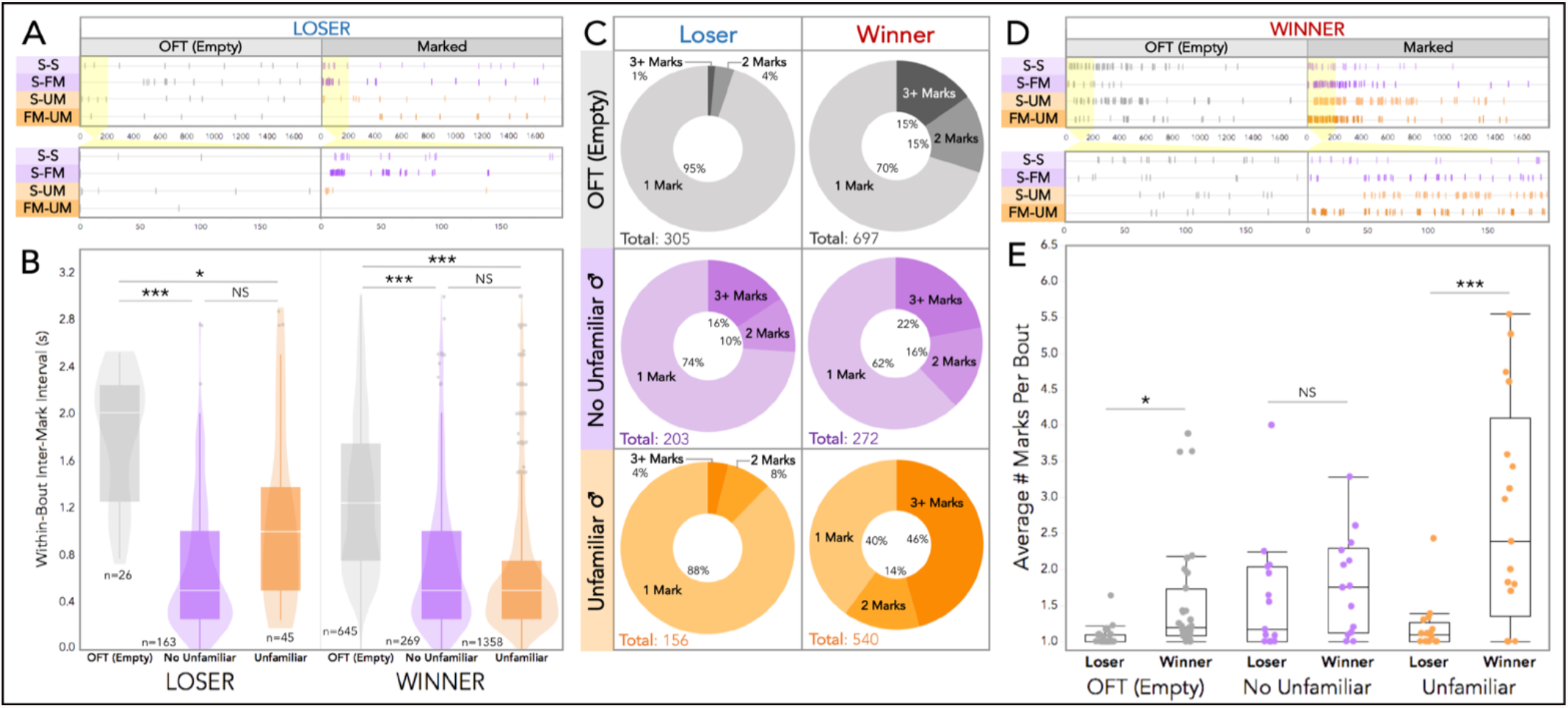
Temporal dynamics of urine signal allocation across scent-marked contexts. **(A)** Example data event plots depicting urine marking in OFT and Marked trials of four losing males, each exposed to one of the four different scent-marked treatments: self-self (S-S), self-familiar male (S-FM), self-unfamiliar male (S-UM), and familiar male-unfamiliar male (FM-UM). The event plot for the entire trial duration is shown on top and a zoomed-in view of the first 200s is shown below. **(B**,**C**,**D)** Urine-marked treatments are grouped as “no unfamiliar male” urine (purple: S-S and S-FM) and “unfamiliar male” urine (orange: S-UM and FM-UM). **(B)** Box and violin plots of within-bout IMIs by fight outcome and trial group: OFT, No Unfamiliar (S-S & S-FM), and Unfamiliar (S-UM & FM-UM). **(C)** Donut plots by trial and trial group depicting the proportion of bouts composed of: 1 mark, 2 marks or 3+ marks. Mark totals in the bottom left-hand corner. **(D)** Example data event plots depicting urine marking in OFT and Marked trials of four winning males, each exposed to one of the four different scent-marked treatments: S-S, S-FM, S-UM, FM-UM. The event plot for the entire trial duration is shown on top and a zoomed-in view of the first 200s is shown below. **(E)** Boxplot of the average number of marks per bout by fight outcome and trial group. **(B**,**E)** Linear mixed models (LMMs) were used to model relationships, analyses of variance (ANOVAs) were used to test for overall effects, and post hoc pairwise comparisons were performed using the *emmeans* package (significance codes: NS p>0.05, * p<0.05; ** p<0.01, *** p<0.001).

The number of marks per bout changes with social outcome and scent-mark type (Figure 5C). The average number of marks deposited per bout is significantly shaped by scent-mark familiarity (*F*_1,76_ = 13, *p* = 1.3e-05) and fight outcome (*F*_1,66_ = 22, *p* = 1.9e-05), with a strong two-way interaction (*F*_1,76_ = 6.3, *p* = 0.0031; Figure 5E). Fight outcome and familiarity both influence the composition of marking bouts (Figures 5C & 5E). In an environment empty of scent marks (OFT), winners allocate considerably more multi-mark bouts than losers (30% vs 5%; Figure 5C). And the average number of marks per bout is significantly higher among winners (*t*_1,105_ = -3.0, *p* = 0.027; Figure 5E). Interestingly, the differences in bout composition narrows in scent-marked trials with familiar-only urine (Figure 5C). In these trials, winners deposit slightly more multi-mark bouts (38%), while losers dramatically shift the amount of multi-mark bouts (26%; Figure 5C). The average number of marks per bout similarly does not differ between winners and losers in familiar-only trials (*t*_1,117_ = -1.0, *p* = 0.87; Figure 5E). The reverse is true for trials with unfamiliar male urine present (Figure 5C). Here, losers produce bouts with similar bout compositions to the empty OFTs (Figure 5C). Winners double the proportion of multi-mark bouts compared to the OFTs (60%), and many of these bouts contain at least 3 marks (46%; Figure 5C). Expectedly, the average number of marks per bout is significantly higher among winners when unfamiliar urine is present (*t*_1,117_ = -5.6, *p* = <0.0001; Figure 5E). Thus, the dynamic temporal patterns of urine allocation change in response to the identity of urine to countermark.

## Discussion

Taking advantage of a novel thermal imaging approach, we uncovered dramatic shifts not only in the amount of urine marking but also the relative timing of urine deposition in response to varying social environments and recent social experiences. In contrast to the dynamic changes observed for some features of scent marking, we found that initial marking strategy significantly explained allocation efforts days later. Our study reveals a mixture of static and dynamic features of urine marking behavior in response to different social contexts. At the start of trials, all males were fully adult and sexually mature with similar prior experiences. Nevertheless, we found substantial variation in the amount, latency, and timing of scent marking in the first trial (Figures 2 & 3). Competitive social experiences had multiple effects on scent marking behavior including priming effects on scent mark timing as well as winner and loser effects. The temporal rhythms of scent marking behavior changed over the course of the trials and indicate a dynamic and socially responsive feature of signaling behavior that had been previously unobserved.

Winning or losing an aggressive contest has strong and long-lasting effects on signaling decisions (Figures 4 & 5), consistent with classically described winner-loser effects (Dugatkin, 1997; Harrison et al., 2018; Hsu & Wolf, 1999). These winner-loser effects most prominently alter total allocation effort and marking bout composition. As described in the literature, we find males quickly downregulate urine allocation after losing a competitive contest (Arakawa, Blanchard, et al., 2008; Desjardins et al., 1973). Though signal allocation is influenced by recent social outcomes, we find that initial signal investment has both stable and robust effects on marking behavior. In other words, where males start off influences their signaling decisions days later. Low-marking individuals remain relatively low-marking, and high-marking individuals stay relatively high-marking. The magnitude of the observed winner-loser effects is therefore contingent on the initial investment decisions of males.

We detect “silent” low-marking winners in our dataset, for which we could find no previous description in the literature. It’s noteworthy that studies do sometimes pre-screen males for baseline urine marking behavior in scent-marking assays, which we did not do (Kaur et al., 2014). The detection of multiple apparently competitive males that mark at very low rates suggests several possible hypotheses to explain their low marking behavior. First, the result may be driven in part by our trial design. By pairing evenly-matched males, it could be we observed contests in which low-marking males won a fight because two relatively low-marking individuals were paired together. Additionally, better-than-expected outcomes could give rise to slower response times than is observed for worse-than-expected outcomes, in which high-marking losers rapidly downregulate signal allocation (Arakawa, Blanchard, et al., 2008; Desjardins et al., 1973). Second, the low-marking males may differ in some aspect of hydration physiology that we did not measure. Species and strains of mice vary in water intake and urination levels (Bittner et al., 2021; Fertig & Edmonds, 1969; Moro & Bradshaw, 1999), though we observed low marking winner males from both strains used in this study. Third, being a “silent” yet competitive male might represent a distinct signaling strategy in house mice. Given the high metabolic costs of signaling, it’s plausible that some males withhold signal investment to continue investing in body mass or to avoid detection by other males. House mice males may pursue diverse strategies, including the classically described “territorial male” that invests highly in urine marking and territory as well as scent-silent “sneaker males” (Aubin-Horth & Dodson, 2004; Bhandiwad et al., 2017; Miles et al., 2007; Sinervo & Lively, 1996; Zamudio & Sinervo, 2000). While our data does not directly test this relationship, the frequency of low-marking winners warrants further investigation. Certainly, the simple correlation between marking and dominance is considerably more complex than previously described, and the interaction between competitive social experience and initial investment is crucial to understanding male signaling decisions.

Several features of marking behavior are primed by competitive experience. Mice mark more rapidly after a contest, regardless of outcome (Figure 2). The time between deposition events similarly shrinks, such that marking bouts transition from chain-line mark sequences to rapid bursts, resulting in fundamental shifts in motor pattern sequences. Aggressive contests thus appear to push males into a competitive state, driving changes in fine motor adjustments in marking behavior. Voluntary, involuntary, and context-dependent urination are all mediated by neuronal subpopulations in Barrington’s nucleus in the brainstem (Hou et al., 2016; Keller et al., 2018; Verstegen et al., 2019). The fine-scale modulation of urinary motor control we observe in our dataset adds additional complexity to this underlying circuitry, as our data reveal social interactions modulate the rhythms of urination behavior. This result opens the possibility for future work to examine how social experience influences finely-controlled motor outputs in an important social signal. Particularly as in more natural environments the rhythm and timing of urine deposition may be crucial for efficient signal allocation as males traverse large distances and defend territories.

Males show markedly different responses to urine based on their familiarity with the scent and their recent fight outcome. Past studies have shown that males can finely distinguish self from non-self urine (Kaur et al., 2014) and that female mice recognize specific males based on their urine marks (Hurst et al., 2001, 2005). We had designed trials with scent-marked corners with urine of different males expecting to see evidence that males allocated marks differently to each corner. We did not detect clear effects of differential allocation to marked corners, suggesting two non-mutually exclusive possibilities: (1) the scent of unfamiliar males may be much more important in driving allocation decisions and (2) the size of the arena may be too small for the marks to be in appreciably different locations from the perspective of the mouse. Established competitive relationships, or familiarity, with another male profoundly shapes marking behavior. Familiarity affects most facets of urine marking, including: total allocation, timing of deposition, and marking bout composition. Surprisingly, losers tend to increase mark allocation effort and display more frequent bursts of multi-mark bouts in response to familiar scent marks. In contrast, winners downregulate their marking efforts toward familiar urine marks. Male mice are known to increase countermarking toward non-self male urine (Hurst & Beynon, 2004; Kaur et al., 2014); however, we find that this effect is strongly modulated by recent experience (or lack thereof) with the urine’s producer. We find evidence for the dear enemy effect in winners and the nasty neighbor effect in losers, suggesting that recent social experiences modulate how animals invest in territorial advertisement and signaling. The dear enemy effect has been reported in a wide array of species (Booksmythe et al., 2010; Briefer et al., 2008; Christensen & Radford, 2018; Siracusa et al., 2021; Tumulty & Bee, 2021; Zorzal et al., 2021). In our study, the responses toward familiar males are even more stark when compared to how males respond to unfamiliar urine marks. Winners dramatically upregulate all competitive marking efforts and losers essentially go completely scent “silent.” Males remain aggressive and vigilant toward unfamiliar or new neighboring males under the dear enemy model, and winners in our trials aggressively mark toward novel males as well. Novel males would represent a threat to their current dominance status, motivating winning males to invest further in signaling behaviors. Losers on the other hand are at risk of an additional aggressive encounter after being recently defeated in a contest. By staying “silent” losers may avoid further conflict with a new territorial contender, potentially in a “wait-and-see” strategy (Olivier et al., 1991; Rychlik & Zwolak, 2005). However, when presented with urine of the male that recently defeated them, losers actually upregulate marking efforts. This response may be a “nasty neighbor” effect, in which the threat of familiar territorial males exceeds that of strangers (Christensen & Radford, 2018). Alternatively, this increased marking response in losers could be a form of subordinate marking to mediate recognition and thereby reduce aggression with a familiar competitor. Male house mice thus actively track multiple identities in the environment and dynamically adjust their signaling decisions in terms of total allocation and timing.

This work underlines the importance of examining signal responses across variable social odor landscapes and understanding the decision rules behind costly and complex behaviors. Because thermal recording allows for high-throughput and detailed analysis of scent marking, future studies can explore even more detailed social contexts and odor environments. This is particularly relevant given that our results provide new evidence for possible alternate signaling strategies, as well as complex territorial dynamics reflecting dear enemy and/or nasty neighbor effects in house mice. Lastly, the implementation of this novel thermal recording method has the potential to reveal important features underlying the neurophysiological basis of socially-modulated and voluntary urination behaviors.

## Materials and methods

### Key resources table

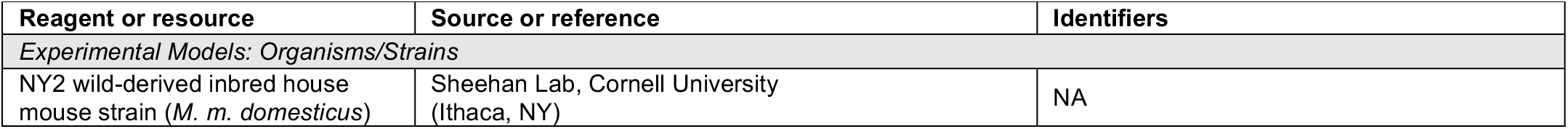
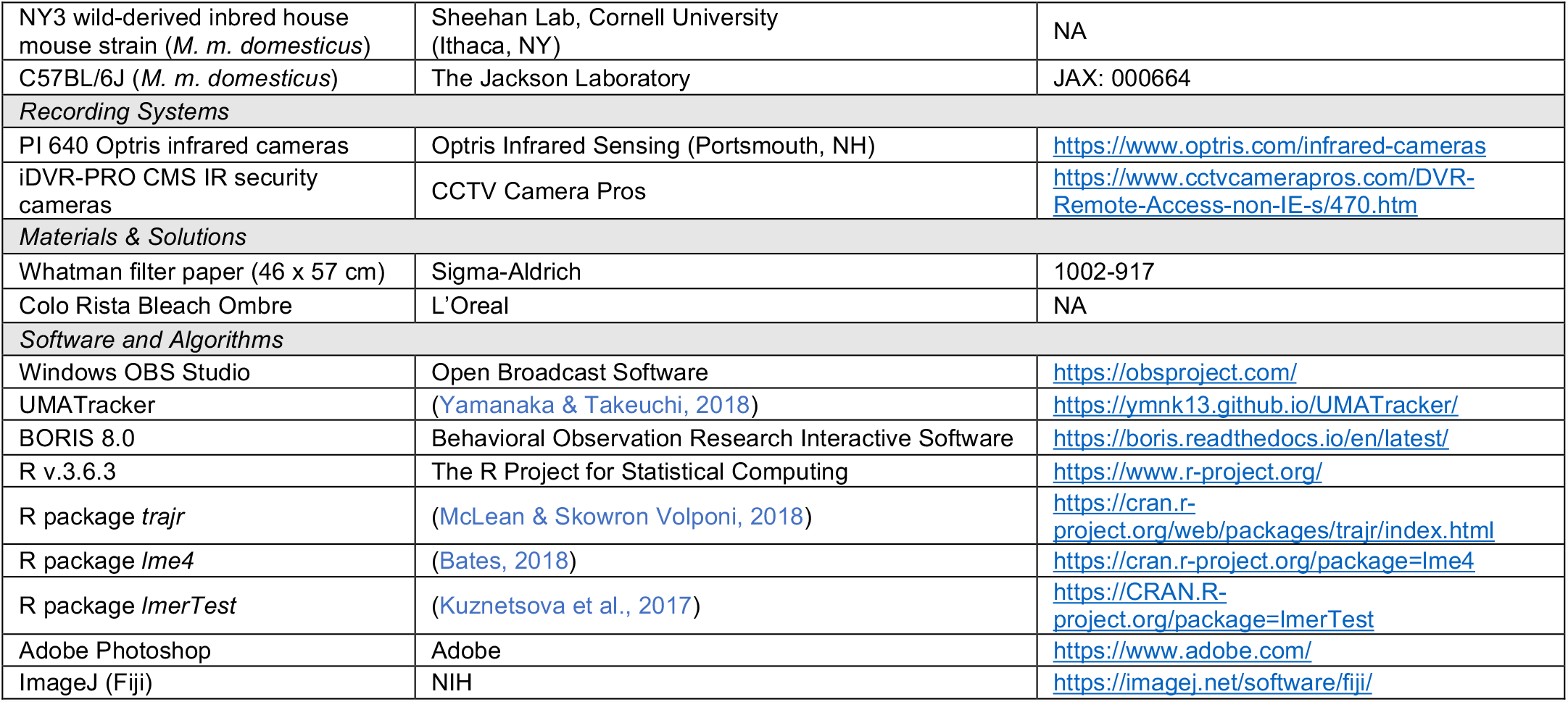

#### Animals

All experimental subjects in this study were males (n=62) from two wild-derived partially inbred strains (NY2 and NY3) of house mice (*Mus musculus domesticus*). Parental generations of these strains were caught in Saratoga Springs, NY by M. Sheehan (Phifer-Rixey et al., 2018). Wild-derived strains were used because competitive behaviors characteristic of wild mice are less pronounced in highly inbred and domesticated laboratory strains (Chalfin et al., 2014; Tuttle et al., 2018) and inbred strains tend to share identical urinary protein profiles (Cheetham et al., 2009). At weaning age (3-4 weeks) males were placed into a holding cage alone for 1-2 weeks, and were subsequently paired with a female to allow for sexual experience as sexually naïve mice are known to exhibit different social behavior (Stowers & Liberles, 2016). All males were allowed to reach adulthood, were between 3-5 months old by the time of experimental testing, and had the opportunity to produce one or more litters. All holding and breeding cages contained corn cob bedding, cardboard huts, and cotton nestlets. Mice were maintained in an Animal Care facility at Cornell University with a 14:10 shifted light:dark cycle (lights went out at 10 PM and on at 12 PM) and were provided food and water *ad libitum*. Mice were handled minimally and with transfer cups whenever possible to reduce stressful handling.

#### Urine collection

Urine was collected from each male subject and from C57BL/6 males to present self, familiar male (paired competitor), and unfamiliar male (C57BL/6) urine in the urine-marked zones on the final day of the trial series (Figure 1A). Urine collection was performed using the single animal method: males were placed atop a metal grate (an upside down cage hopper) over a clear plastic bag for 30 minutes to 1 hour (Kurien & Scofield, 1999). Males were subsequently taken off the plastic bag and returned to their breeding cage. The urine droplets present on the plastic bag were collected and stored at -80°C until use. Urine collected from subject males was stored individually until the day of the urine-marked trials (Day 4: Figure 1A). For sufficient urine volume for the urine-marked trial treatments (Figure 1A), between 200-400uL was collected from each NY2 and NY3 male subject. On the day of the urine-marked trials, individual aliquots for a subject male were thawed on ice and pooled together. Urine collection for the C57BL/6 males was performed the month prior to beginning experimentation. We collected at least 50uL of urine each from over 20 adult breeding C57BL/6 males, urine was stored on the day of collection at -80°C. Once a sufficient volume was collected, individual aliquots were thawed on ice, and all C57BL/6 male urine was pooled into a single volume and subsequently aliquoted and stored at -80°C. This was done such that all unfamiliar C57BL/6 urine stimuli presented to males across trials were as similar as possible.

#### Behavioral experiments

All handling was performed with transfer cups throughout the duration of the trials to minimize stressful handling confounds. One day prior to experimentation, we recorded subject male body weights to size-match individuals as closely as possible (average weight difference: 2.4g). All males were in breeding cages at the time of the experiment and most successfully reproduced (84%) prior to the trial series. As house mice are nocturnal, all experiments were conducted in the dark during the dark cycle to ensure ethological accuracy (Peirson et al., 2018). All experimentation occurred between 12 PM and 4 PM to minimize circadian variation. Additionally, experimentation concluded prior to the winter months. Laboratory mice exhibit seasonal variation with respect to certain physiological parameters like serum concentrations of sex hormones, suggesting a possible mechanism for the internalization of annual time independent of light cycle, temperature, and humidity (Mock et al., 1975). While the available literature provides conflicting evidence as to whether these effects extend to behavior, we nonetheless took measures to avoid such confounds (Ferguson & Maier, 2013). Trial series were performed in sets of between 2-5 male pairs.

Behavioral trials consisted of a 4-day trial design, in which age and weight-matched adult breeding males of distinct wild-derived strains (NY2 and NY3) were paired as competitors and presented a series of social and scent-marked trials (Figure 1A). We pair-matched each NY2 mouse with a NY3 mouse to ensure that no two paired mice were genotypically identical and that their scent marks were perceptibly different (unique major urinary protein profiles) (Kurien & Scofield, 1999), resulting in a total of 31 pairs (n=62). To ensure identification of males within a pair (NY2 and NY3 strains are visibly indistinguishable), we ear-clipped and bleached a patch of rump fur of one male in each pair a week prior to experimentation. Mice were anesthetized with isoflurane (5%). A heating pad was used to maintain a stable body temperature. Isoflurane was delivered at 1-3% throughout the bleaching procedure. L’Oreal Colo Rista Bleach Ombre (salon bleach) was mixed as per the manufacturer’s instructions and dabbed onto the top layer of fur using a sterile cotton swab. Care was taken to prevent bleach from contacting the skin. Twenty minutes after application, sterile cotton tipped swabs dipped in water were used to rinse the bleach from the fur. The fur was then dabbed dry with paper towels. Mice were placed under a heat lamp for 5 minutes or until they were fully recovered from anesthesia before being transferred back to their home cage.

All trials were performed in one of two trial chambers that were sound proofed and fitted with recording systems. For all trials large sheets of Whatman filter paper lined the floor of each trial to collect urine blots and to present urine stimuli. The same size PVC arenas were used throughout (50 cm x 50 cm), though split in half with the mesh barrier for the Mesh trials (Figure 1A). At the end of each trial, males were placed back into their breeding cages. On Day 1 of the trial series, paired males were placed on either side of a wire mesh barrier in an arena for 30 minutes (Mesh 1, Figure 1A). At the end of the 30 minutes, males were briefly removed from the arena into large transfer cups, the filter paper was labeled and removed, a fresh filter paper was placed in the arena, and the mesh barrier was removed. Males were placed back into the arena for a 30 minute aggressive contest (Fight trial, Figure 1A). On Day 2, each male was placed alone in an empty “Open Field” arena without any stimuli aside from the arena itself for 30 minutes (Open Field trial, Figure 1A). On Day 3, males were placed back into the Mesh arena for 30 minutes with the same male competitor encountered on the first day, without the subsequent fight trial (Mesh 2 trial, Figure 1A). On Day 4, males were placed into the arena alone and subjected to a 30 minute urine-marked stimulus trial, consisting of one of 4 possible treatment types. Each treatment included two spatially distinct urine-marked zones placed in opposite corners of the arena (front right – back left vs. back right – front left). Urine-marked corner zones contained aliquoted male urine of 3 possible identities: self, familiar, or unfamiliar male. Self-urine was collected from the focal trial mouse; familiar male urine was from the paired male competitor of the focal mouse; unfamiliar male urine was from pooled C57BL/6 urine. The urine stimuli for a urine-marked trial was thawed, pooled and kept on ice until aliquoted for the urine-marked stimulus trial onto filter paper. Urine stimuli were placed on the filter paper directly before the trial start in standardized locations and volumes. The four treatment types span a range of scent mark combinations: self-self, self-familiar, self-unfamiliar, familiar-unfamiliar. Paired males (winner-loser pairs) received the same urine-marked stimulus treatment. For all scent marked trials (Days 1-4) the first and last minute of each trial was trimmed prior to analysis. This was done to minimize detection of startle-based urination events caused by placement of mice into arenas and any jostling caused during trial set-up and take-down. The total analyzed trial length was thus 28 minutes.

Trials and treatments were randomized as follows. Male trial order and arena chamber was pseudo-randomized each day to avoid confounds in arena location and marking behavior over the course of the designated trial period (12 PM – 4 PM). The orientation within the Mesh 1 trials was also randomized (whether males were placed near the back or front of the arena) to account for variation in sound disturbances for males closer to the chamber door; orientations were subsequently flipped for each pair in Mesh 2. Urine-marked trial treatments were pseudo-randomly assigned to each male pair, to ensure similar numbers of male pairs were exposed to the 4 treatment types across sets of trials series. The orientation of urine stimuli was randomly assigned to corner orientations (front right – back left vs. back right – front left). Lastly, the fur bleaching for male identification was performed on one mouse strain (NY2 or NY3) for each trial set, but the bleached strain was switched between trial sets to prevent errors within a trial set and to avoid bleaching only one strain across trial sets.

#### Recording methods

All trials were recorded with a security camera system (iDVR-PRO CMS) at 1080p and 30 frames per second to visualize the high-speed aggressive encounters and to clearly distinguish the male identities (ear-marked and bleached fur). All trials (including fight trials) were recorded thermally using an infrared camera system (PI 640; Optris Infrared Sensing). Thermal cameras were fitted with 33° x 25° lenses and mounted above the experimental arena chambers such that field-of-view for each camera covered the entire arena. The thermal detection window was set at: 61°F - 107°F. Data frames were collected at the max speed, averaging at 3 Hz. Thermal video data was saved by screen-capturing live Optris video output using OBS Studio software. Raw temperature data was also collected in semicolon-delimited CSVs, providing a readout of the temperature in each pixel for each frame.

#### Behavioral scoring

All videos were scored using Behavioral Observation Research Interactive Software (BORIS) (Friard & Gamba, 2016). For the fight trials (Figure 1A), we scored the following aggressive behaviors: chasing, hitting, boxing, and wrestling bouts (Figure S1C) using the infrared security camera video recordings. To score urine mark deposition events Optris thermal video recordings were used for all trials. Urine depositions were scored as a clear hot spot following the focal mouse’s trajectory that subsequently cooled below substrate temperature. Fecal depositions could be eliminated as they are frequently cooler upon deposition event, cool much more slowly, have a distinct shape, and are typically moved around the arena quickly. In addition to scoring the timing of urine deposition events, the placement of urine marks was also scored. Using screen annotation software, we drew precise lines on the video observation corresponding to regions of interest for each thermally-recorded trial (Figure 1B-C).

#### Tracking

Mice were tracked using the software UMATracker (Release 12) (Yamanaka & Takeuchi, 2018). Infrared security camera recordings were used to track focal mouse movement, as the video were recorded at a higher framerate. Filters were generated using the following modular settings (in order): output – Closing: Kernel = 6 – Opening: Kernel = 6 – Threshold: 100 – BGRToGray – input. Videos were tracked using Group Tracker GMM algorithm. Area51 was used to generate desired regions of interest for each trial (Figure 1B-C) and analyze the relative space use in each of these regions. The R package *trajr* (McLean & Skowron Volponi, 2018) was used to quantitatively characterize the following information from the tracked data frames: speed, acceleration, and trajectory length.

#### Urine blot imaging and processing

Trials were run on Whatman filter paper substrate. Arena edges were outlined with pencil on the filter paper at the end of each trial. We collected all sheets of filter paper used in experimentation (except for the Fight trial) and photographed them under ultraviolet (UV) light. We used three UV bulbs to evenly distribute light on the large filter paper area. Images were converted to greyscale in Adobe Photoshop and the magentas were reduced to ∼20% to observe edges of urine marks clearly. Greyscale images were subsequently processed in ImageJ (Fiji). We subtracted background pixels for a cleaner image (100 px), applied image thresholding (manually adjusted when necessary), and converted images to binary in order to convert to mask, fill holes and perform watershed algorithm. This processed image was then used to analyze the number of particles, with Size (pixel^2): 100-Infinity and Circularity (0-1.00).

#### Urine mark bout classification

The median inter-mark interval (2.99 seconds) for all males across all trials was used to determine whether marks get clustered into a marking “bout” (Figure S4A). Any two marks that occur in sequence with an inter-mark interval less than 3 seconds are clustered together into a multi-mark bout, allowing us to examine within-bout temporal dynamics.

#### Statistical analysis

We conducted all statistical analyses in R 3.6.0 (R Development Core Team 2019). We used linear mixed models (LMMs) and paired statistical tests to examine relationships between dependent and response variables. Models were fitted using the package *lme4* (Bates, 2018). The *lmerTest* package was used to calculate degrees of freedom (Satterthwaite’s method) and p-values (Kuznetsova et al., 2017). Dependent variables were transformed for a subset of models to meet assumptions for model residuals after visually inspecting model residuals. We used a type 3 analysis of variance (ANOVA) to test for overall effects of fixed factors or interactions in the models. Post hoc comparisons were conducted using the *emmeans* package (Lenth, 2016). R script and data sheets used for all statistical analyses are provided.

## Acknowledgements

We thank Kevin Besler, Christen Rivera-Erick and Melanie Colvin for crucial technical assistance; Russell Ligon and Caleb Vogt for helping establish recording systems and tracking methods in the lab; and James Tumulty for manuscript feedback.

## Additional information

### Funding

| Funder | Grant reference number | Author |
| --- | --- | --- |
| USDA Hatch Grant | NYC-191428 | Michael J Sheehan |

The funders were not involved in the design of the study; the collection, analysis and interpretation of data; the writing of the manuscript and any decision concerning the publication of the paper.

### Ethics

All experimental protocols conducted at Cornell University were approved by the Institutional Animal Care and Use Committee (IACUC: Protocol #2015-0060) and were in compliance with the NIH Guide for Care and Use of Animals.

### Authors’ contributions

CHM and MJS conceived the study. CHM performed trials and analyses. MFH, JY, BCC, KH and AYL collected samples, scored behavioral trials, and generated tracking data. CHM and MJS wrote the initial drafts of the paper. MRW edited the manuscript. All authors contributed to manuscript preparation.

### Competing interests

The authors declare no competing interests.

## Additional files

### Supplementary figures

**Figure S1.**
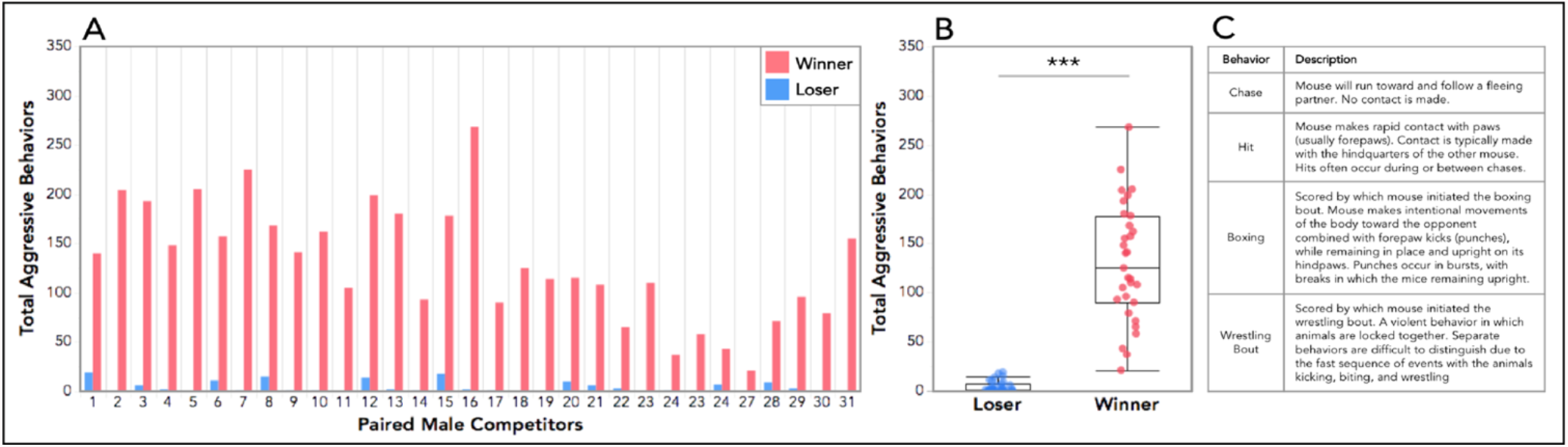
Male aggressive behaviors scored in contests (fight trials) between paired competitors. **(A)** Total aggressive behaviors performed by each paired male competitor. The fight outcome (the categorization of winners and losers) was determined by which male performed more aggressive behaviors within a pair. **(B)** Across all 31 pairs, winning males performed significantly more aggressive behaviors than losing males (*t*_*1,31*_= -12.6, *p* = 1.09e-13). Welch’s t-test was used to compare the total aggressive behaviors performed by the two fight outcome categories (significance code: *** p<0.001). **(C)** Ethogram used to score aggressive behaviors. State events: chase, boxing and wrestling bouts. Points events: hits. An event is only coded for a male subject if the individual initiated the behavior (i.e. wrestling bout is coded for only one participant – the initiator – of that event).

**Figure S2.**
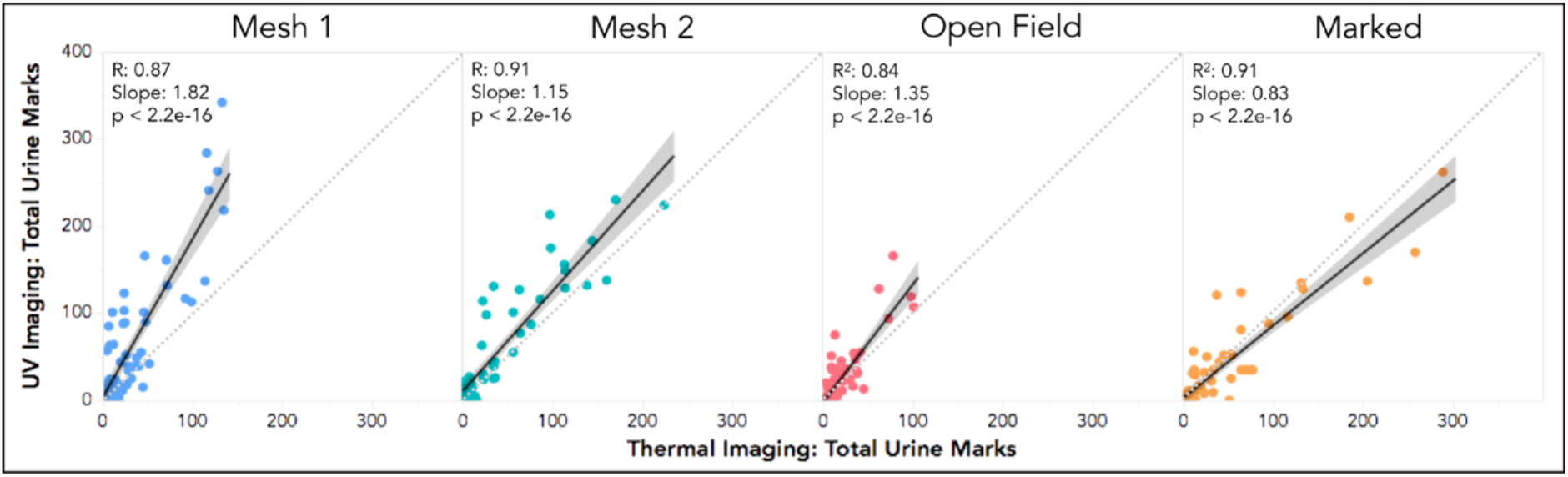
Comparison of urine mark detection methods across trial types: Ultraviolet light (UV) blot imaging vs. thermal imaging. The two detection methods are well-correlated with each other (R > 0.8). For both Mesh trials and the Open Field trials, UV imaging consistently detected more urine marks than thermal imaging. The Marked trials revealed the opposite pattern, with thermal imagining detecting more urine marks than UV imaging. Three trials were excluded from this dataset due to poor urine blot quality, and one trial was excluded as an outlier.

**Figure S3.**
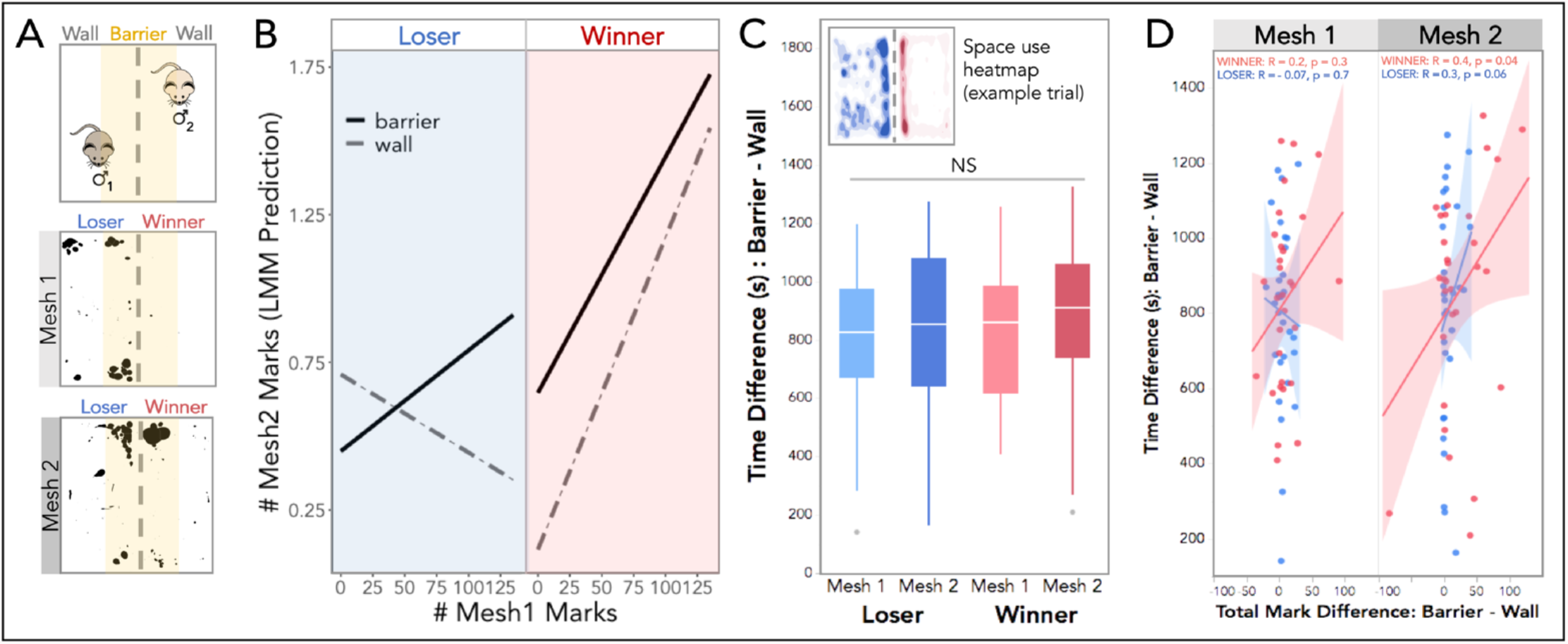
Mesh trial spatial marking and space use. **(A)** Top: schematic of the mesh trials indicating the social “Barrier” (yellow) and non-social “Wall” regions of interest (ROIs). Below: Example urine blots of a male pair (winner and loser) pre- and post-fight demonstrating the spatial allocation of urine marks at the social boundary. **(B)** Linear mixed model (LMM) prediction of the total number of marks in the post-fight mesh trial (Mesh 2) given the fight outcome (winner=red; loser=blue), initial signal investment (# Mesh 1 marks), and the ROI (barrier=solid; wall=dashed). **(C)** Difference in time (s) spent in the Barrier vs. Wall regions of interest (ROIs) across mesh trials by winning and losing males. Winners and losers spend more time at the social boundary (Barrier) across mesh trials. Top left corner: an example heat map a male pair in a mesh trial (Mesh 1), depicting how all males spend more time at the social boundary (Barrier) than the non-social ROI (Wall) across mesh trials, regardless of fight outcome. **(D)** Comparison of the difference in time spent vs. the difference in total marks allocated in the ROIs (Barrier – Wall) by winners and loser across trials. In both mesh trials, space use and changes in urine allocation effort are not detectably correlated with each other among winning or losing males (R < 0.2). **(B-D)** Linear mixed models (LMMs) were used to model relationships, analyses of variance (ANOVAs) were used to test for overall effects, and post hoc pairwise comparisons were performed using the *emmeans* package (significance codes: NS p>0.05, * p<0.05; ** p<0.01, *** p<0.001).

**Figure S4.**
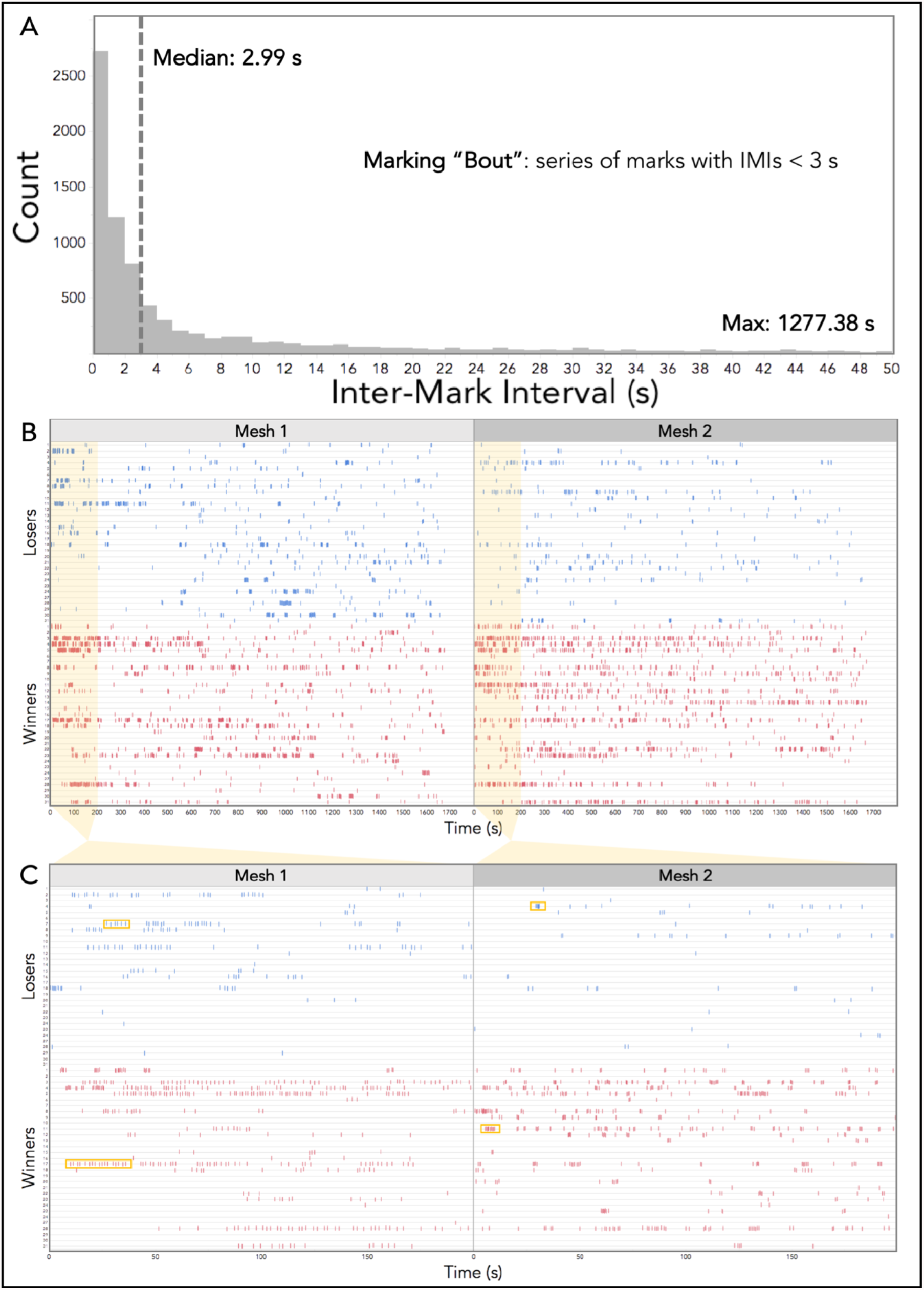
**(A)** Histogram of the inter-mark intervals (IMIs) for all males across all trials. The median value is indicated with a dashed line (2.99 seconds). The range of IMIs extends to nearly the full trial length (only the first 50s is shown), The maximum values are reported in the bottom right corner. The median IMI value was used to define a marking “bout.” Such that any two marks that occur in sequence with an IMI < 3 seconds are grouped together into a multi-mark bout. **(B)** Event plots depicting the urine marking of all male competitors over the course of both mesh trials (Mesh 1=left, Mesh 2 = right) for the entire trial duration (1800 seconds). Pair IDs are indicated on the left-hand axis. Losers depicted on top in blue, and winners on the bottom in red. **(C)** Event plots depicted a zoomed-in view of the first 200 seconds of the trials for all individuals. Example chain-like bouts are outlined in the Mesh 1 panel, and example burst-like marking bouts are highlighted in the Mesh 2 panel (yellow boxes).

**Figure S5.**
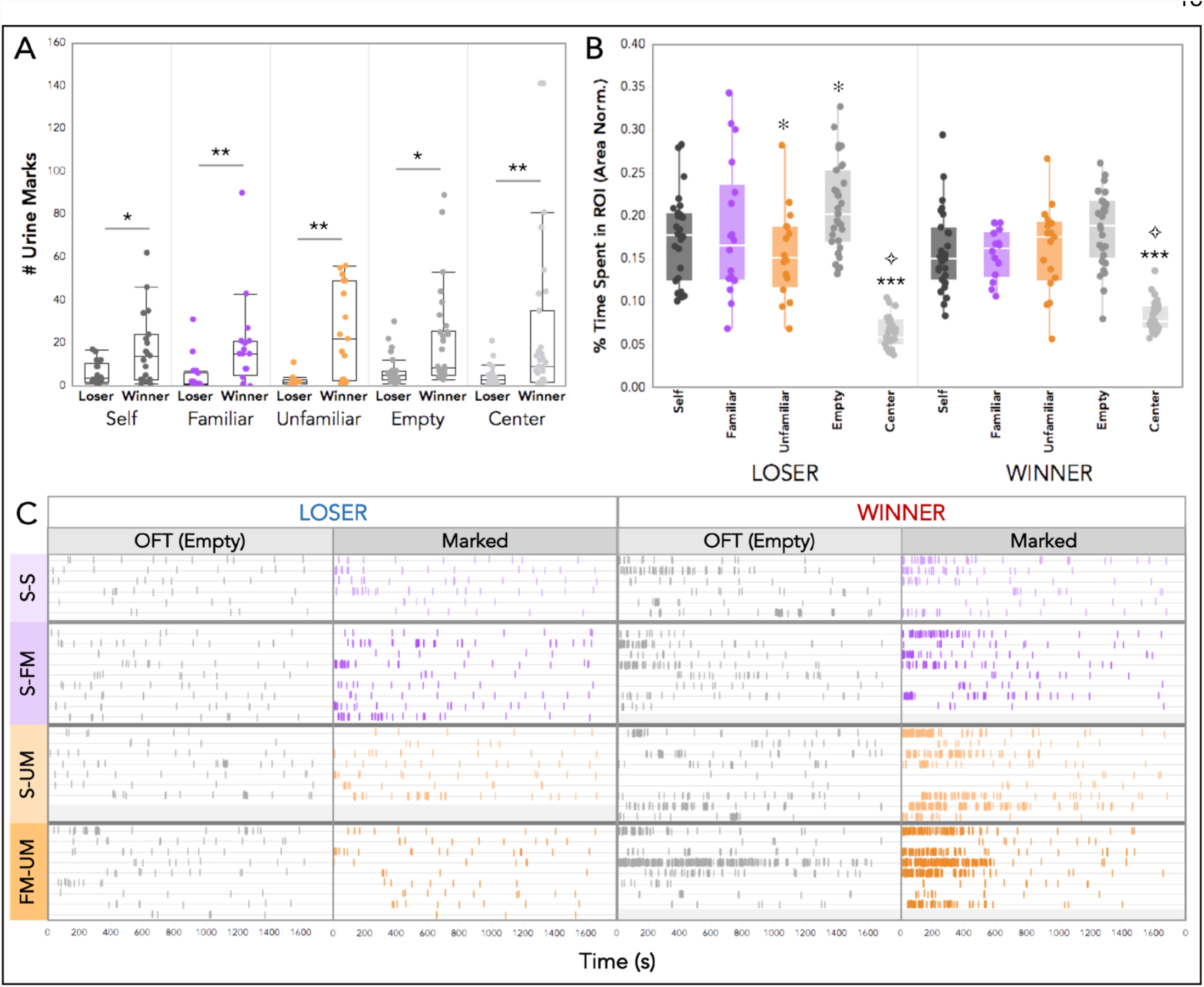
**(A)** Total number of marks deposited by winners and losers in scent-marked trials to a specific ROIs: scent-marked corners (containing self, familiar, or unfamiliar male urine), empty corners, or the center of the arena (significance codes: * p<0.05; ** p<0.01). **(B)** The percent of time spent in by winners and losers in specific urine-marked trial ROIs, normalized to the total area of each ROI (to account for the center being a larger area). Winner and losers spend significantly less time in the Center ROI than all corner ROIs (Self, Familiar, Unfamiliar or Empty; *** p<0.001). Losers spend significantly less time in the Center ROI than winners (✧ p<0.01). Losers spend significantly less time in Unfamiliar corners relative to Empty ones (* p<0.05). **(A**,**B)** Linear mixed models (LMMs) were used to model relationships, analyses of variance (ANOVAs) were used to test for overall effects, and post hoc pairwise comparisons were performed using the *emmeans* package. **(C)** Event plots depicting the urine marking of winning and losing males to the OFTs and the urine-marked trials for the entire trial duration (1800 seconds). Males are grouped by the four different scent-marked treatments: self-self (S-S: light purple), self-familiar male (S-FM: dark purple), self-unfamiliar male (S-UM: light orange), and familiar male-unfamiliar male (FM-UM: dark orange). Almost all male pairs experienced the same treatment, three pairs received different urine-marked treatments due to urine stimuli collection constraints (hence some of the treatment groups have unequal paired number across fight outcome groupings).

## Notes

### Competing Interest Statement

The authors have declared no competing interest.

